# Sister chromatid repair maintains genomic integrity during meiosis in *Caenorhabditis elegans*

**DOI:** 10.1101/2020.07.22.216143

**Authors:** Erik Toraason, Cordell Clark, Anna Horacek, Marissa L. Glover, Alina Salagean, Diana E. Libuda

## Abstract

During meiosis, the maintenance of genome integrity is critical for generating viable haploid gametes [1]. In meiotic prophase I, double-strand DNA breaks (DSBs) are induced and a subset of these DSBs are repaired as interhomolog crossovers to ensure proper chromosome segregation. DSBs in excess of the permitted number of crossovers must be repaired by other pathways to ensure genome integrity [2]. To determine if the sister chromatid is engaged for meiotic DSB repair during oogenesis, we developed an assay to detect sister chromatid repair events at a defined DSB site during *Caenorhabditis elegans* meiosis. Using this assay, we directly demonstrate that the sister chromatid is available as a meiotic repair template for both crossover and noncrossover recombination, with noncrossovers being the predominant recombination outcome. We additionally find that the sister chromatid is the exclusive recombination partner for DSBs during late meiotic prophase I. Analysis of noncrossover conversion tract sequences reveals that DSBs are processed similarly throughout prophase I and recombination intermediates remain central around the DSB site. Further, we demonstrate that the SMC-5/6 complex is required for long conversion tracts in early prophase I and intersister crossovers during late meiotic prophase I; whereas, the XPF-1 nuclease is required only in late prophase to promote sister chromatid repair. In response to exogenous DNA damage at different stages of meiosis, we find that mutants for SMC-5/6 and XPF-1 have differential effects on progeny viability. Overall, we propose that SMC-5/6 both processes recombination intermediates and promotes sister chromatid repair within meiotic prophase I, while XPF-1 is required as an intersister resolvase only in late prophase I.

## Results and Discussion

### Engagement of the sister chromatid in meiotic DSB repair

During meiotic prophase I, double-strand DNA breaks (DSBs) are induced across the genome [1]. A subset of DSBs must be repaired as interhomolog crossovers to ensure accurate chromosome segregation, and the remaining DSBs are repaired through other mechanisms [3]. Studies in *Saccharomyces cerevisiae* indicate the sister chromatid can be engaged during meiosis [4] and multiple lines of evidence have suggested that the sister chromatid is engaged as a meiotic DSB repair template to repair these remaining DSBs in metazoan meiosis [5–7], but the perfect sequence identity shared between sister chromatids has precluded direct testing of this hypothesis in metazoans.

To determine whether the sister chromatid is engaged as a repair template during *C. elegans* meiosis, we developed a non-allelic sister chromatid repair assay (SCR assay; Figure 1A) that utilizes controlled excision of a Mos1 transposon to induce a single DSB within a genetic reporter that detects repair events using the sister chromatid as a template. Similar to other repair assays in *S. cerevisiae* meiosis [8,9] and mammalian mitosis [10–12] systems, our SCR assay is composed of two tandem reporter sequences. In our assay, the upstream copy encodes a truncated GFP allele driven by a *myo-3* promoter (body wall expression). The downstream copy is driven by a *myo-2* promoter (pharynx expression) and is disrupted with the *Drosophila* Mos1 transposable element [13]. Upon heat shock-induced expression of Mos1 transposase [14], excision of the Mos1 transposon produces a single DSB [5]. Repair of the Mos1-induced DSB via nonallelic intersister or intrachromatid recombination yields restoration of functional GFP sequence and GFP+ progeny. The tissue-specific expression of the resultant functional GFP indicates which specific sister chromatid recombination pathway engaged: 1) an intersister or intrachromatid noncrossover will generate functional *pmyo-2::*GFP expressed in the pharynx; and, 2) an intersister crossover will produce *pmyo-3::*GFP expressed in the body wall muscle. Since allelic recombination will not restore functional GFP sequence, the SCR assay only detects nonallelic recombination outcomes.

**FIGURE 1.**
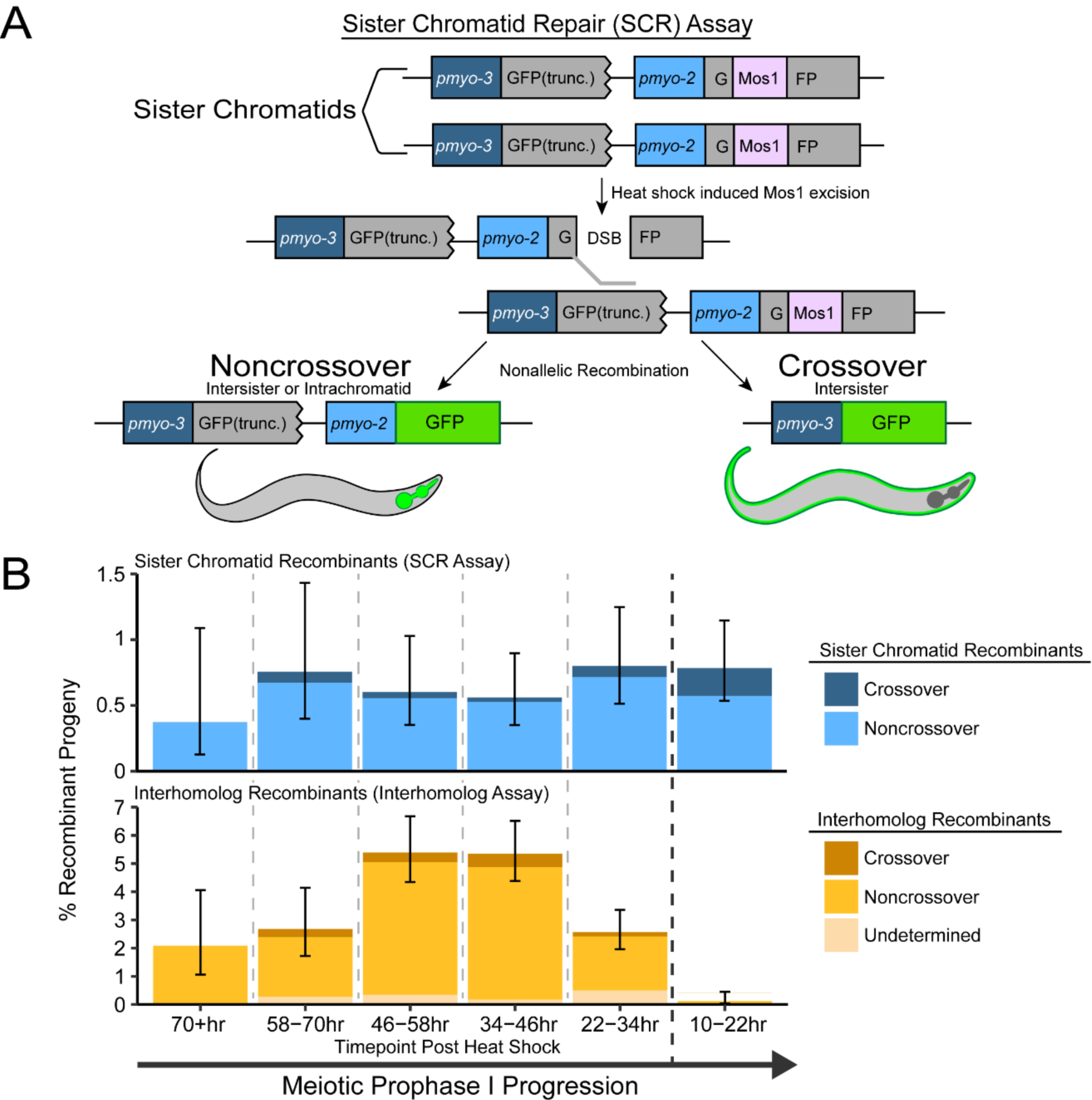
The sister chromatid is engaged as a DSB repair template in meiotic prophase I. (A) Cartoon diagram of the sister chromatid repair (SCR) assay. The SCR assay is composed of two tandem GFP cassettes. The upstream GFP is driven by a *pmyo-3* (body wall) promoter and is truncated, while the downstream GFP is driven by a *pmyo-2* (pharynx) promoter and is interrupted by a Mos1 *Drosophila* transposon. Excision of Mos1 yields a single DSB. Repair of this DSB by intersister or intrachromatid recombination will yield GFP+ progeny. (B) Frequency of recombinant progeny identified in the SCR assay (top) and interhomolog assay (bottom) [5]. Total progeny scored n=SCR assay/Interhomolog assay; 10-22hrs n=3317/1623; 22-34 hrs n=2372/1938; 34-46hrs n=3032/1629; 46-58hrs n=2159/1369; 58-70hrs n=1190/691; 70+hrs n=806/376 (Supplemental Tables 1 and 2). Stacked bar plots represent the overall percent of living progeny that exhibit the indicated recombinant phenotype within a specific time point following heat shock. Error bars represent 95% binomial confidence intervals. Dashed vertical lines delineate between time points scored, while the dark black dashed line delineates between the ‘interhomolog window’ (22-70+hr post heat-shock) and ‘non-interhomolog window’ (10-22hr post heat-shock).

The SCR assay was performed in hermaphrodites heterozygous for the assay at a locus previously assessed for interhomolog repair (exon 6 of *unc-5*; [5]) (see Methods). Since there is no GFP sequence on the homolog in this context, recombination is restricted to sister chromatid and intrachromatid events. With this assay, we observed both noncrossover and crossover GFP+ recombinants at an overall frequency of 0.68% (95% confidence interval (CI) 0.548-0.833%), which represents the frequency of nonallelic recombination at this locus (Figure 1B top). Notably, noncrossovers were the predominant repair outcome from the SCR assay (86.2% of GFP+ recombinants, Supplemental Table 1). This data directly demonstrates that the sister chromatid is engaged as a DSB repair template in *C. elegans* meiosis and enables the assessment of this meiotic DNA repair pathway for the first time in a metazoan.

While the homologous chromosome is the preferred recombination template in meiotic prophase I [3], several studies have hypothesized a switch in template bias from the homolog to the sister chromatid during late meiotic prophase I [5,6,6]. Similar to a previous *C. elegans* assay which assessed interhomolog repair during meiosis (interhomolog repair assay; Figure 1B bottom) [5], the SCR assay can determine the stages of meiotic prophase I in which the sister chromatid can be engaged as a repair template. Given the established timing of meiotic prophase progression for *C. elegans* oogenesis, progeny laid in the 22-70 hour timepoints were derived from oocytes spanning entry into meiotic prophase I through mid-pachytene at the time of heat shock (Mos1 excision), while the oocytes yielding progeny at the 10-22 hour time point were at late pachytene/diplotene [5,16]. While neither the interhomolog assay nor SCR assay detect whether a DSB is repaired within the same meiotic stage it was induced, we can still determine the latest window in which a repair template is available. Specifically, DSBs induced in the interhomolog assay are robustly repaired with the homologous template during the 22+ hour time points (‘interhomolog window’) and not the 10-22 hour time point (‘non-interhomolog window’) (Figure 1B bottom, Supplementary Table 2) [5], the SCR assay demonstrates that DSBs induced at different times throughout meiotic prophase can be repaired using the sister chromatid (or same DNA molecule), and that such repair occurs at similar frequencies regardless of the timing of DSB induction (Figure 1B top, Supplementary Table 1). Thus, while engagement of the homologous chromosome is restricted to a specific window of meiotic prophase I, the sister chromatid may be engaged as a repair template for DSBs induced throughout meiotic prophase I. This data further directly demonstrates that intersister or intrachromatid repair becomes the preferred recombination pathway in late meiotic prophase I when the homologous chromosome is no longer readily engaged for repair (10-22 hours post-heat shock, Figure 1B).

We observed both noncrossover and crossover recombinant progeny at all timepoints except at 70+ hours (Figure 1B top), indicating that DSBs induced throughout meiotic prophase I may be repaired by sister chromatid crossover and noncrossover recombination pathways. Crossover recombinants are specifically enriched in the non-interhomolog window compared to the interhomolog window (Fisher’s Exact Test p=0.037). These results indicate that a late pachytene transition increases DSB resolution by intersister crossover recombination.

### Role of Smc5/6 in sister chromatid repair

The SCR assay enables direct testing of proteins hypothesized to regulate sister chromatid repair [17], such as the conserved structural maintenance of chromosomes 5/6 (Smc5/6) complex [17–19]. To determine if SMC-5/6 promotes sister chromatid recombination in meiosis, we performed the SCR assay in an *smc-5(ok2421)* null mutant. We observed recombinant progeny arising from *smc-5* mutants at timepoints spanning 10-58 hours post-heat shock (Figure 2A, Supplementary Table 3 top). *smc-5* mutants lacked sister chromatid recombinant progeny at the later timepoints (58+ hours post-heat shock), but this absence is due to low sample sizes (Supplementary Table 3 top) [17].

**FIGURE 2.**
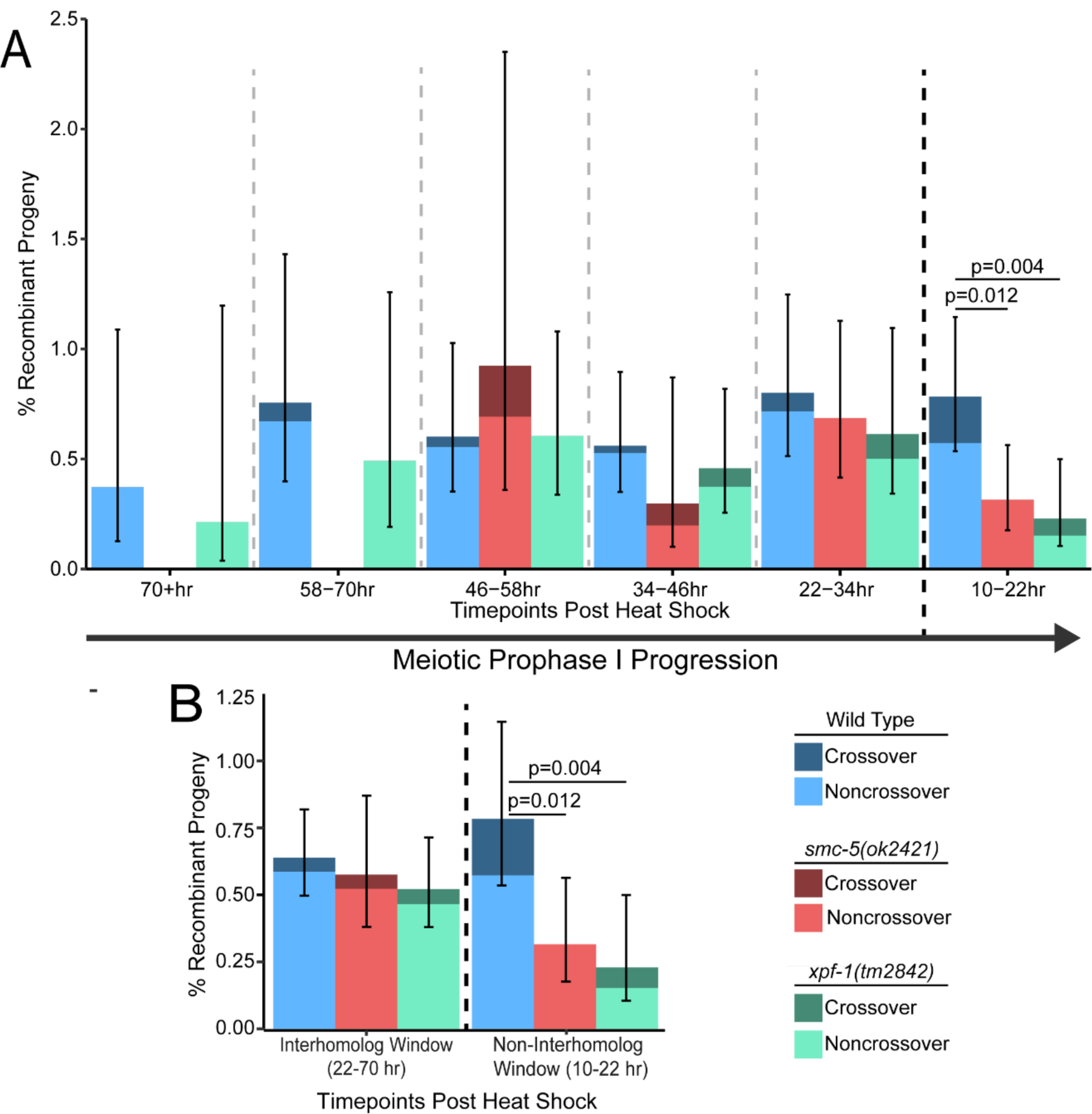
SMC-5/6 and XPF-1 are required for efficient sister chromatid repair in late meiotic prophase I. (A) Frequency of SCR assay recombinant progeny in wild-type, *smc-5(ok2421)*, and *xpf-1(tm2842)* mutants at each scored time point following heat shock. Total progeny scored n=wild-type/*smc-5*/*xpf-1*; 10-22hrs n=3317/3491/2618; 22-34 hrs n=2372/2188/1793; 34-46hrs n=3032/1009/2400; 46-58hrs n=2159/433/1819; 58-70hrs n=1190/121/813; 70+hrs n=806/72/469 (Supplemental Tables 1 and 3). (B) Frequency of recombinant progeny identified in the SCR assay within binned windows of prophase I defined by observation of recombinants in the Interhomolog assay. n=wild-type/*smc-5*/*xpf-1*; interhomolog window n=9559/3801/7294; non-interhomolog window n= 3317/3491/2618; 22-34 hrs (Supplemental Tables 1 and 2). Stacked bars represent the overall percent of living progeny that exhibit the indicated recombinant phenotype within the labeled time interval following heat shock. Error bars represent 95% binomial confidence intervals. P values were calculated by Fisher’s Exact Test. Dashed vertical lines delineate between time points scored, while the dark black dashed line delineates between the ‘interhomolog window’ (22-70+hr post heat-shock) and ‘non-interhomolog window’ (10-22hr post heat-shock).

In the interhomolog window of *smc-5* mutants (22-70hrs, early-mid prophase I), the frequency of recombinants was indistinguishable from wild-type (Figure 2B, Fisher’s Exact Test p>0.05). However, in the non-interhomolog window (10-22 hours, late prophase I), the frequency of observed sister chromatid recombinants in *smc-5* mutants was significantly reduced relative to wild-type (Figure 2A, Fisher’s Exact Test p=0.012). Notably, we observed no crossover recombinant progeny from timepoints 10-34 hours post-heat shock (Figure 2A, expected crossover frequency 20%, Exact Binomial Test p=0.005), indicating that *smc-5* mutants are defective in sister chromatid crossing over specifically in late meiotic prophase I. Together, these data demonstrate that recombination repair may be differentially regulated within meiotic prophase I such that intersister crossover recombination in late prophase switches from SMC-5/6-independent to SMC-5/6-dependent mechanisms.

### Role of XPF-1 nuclease in sister chromatid repair

We next investigated the role of the resolvase XPF-1 in sister chromatid recombination. XPF-1 is the *C. elegans* homolog of the XPF/RAD1 nuclease and acts semi-redundantly with other nucleases to resolve meiotic interhomolog crossovers [20–23]. XPF-1 is also required for single-strand annealing (SSA), a mutagenic homology-directed repair pathway which may be engaged upon exposure of >30bp of repeated sequence on each resected ssDNA strand of a damaged chromosome and results in deletion of sequences between tandem repeats [24–26]. As the SCR assay contains tandem GFP cassettes (Figure 1A), engagement of SSA to resolve Mos1-induced DSBs could yield progeny with a phenotype that could be interpreted as a crossover. To both assess the role of XPF-1 in sister chromatid repair and determine whether our assay is identifying SSA-mediated DSB repair, we performed the SCR assay in an *xpf-1(tm2842)* mutant and observed both noncrossover and crossover recombinant progeny (Figure 2, Supplementary Table 3 bottom).

Neither the overall recombinant frequency nor the proportion of crossover progeny in *xpf-1* mutants differed from wild-type within the interhomolog window (Figure 2B, Fisher’s Exact Test p>0.05). While crossover progeny were not detected at the 34-70+ hour timepoints in *xpf-1* mutants, this absence is likely due to sampling a low incidence population (crossovers) from a limited pool (recombinants) because this deviation was not significant from wild-type (Expected crossover frequency 8%, Exact Binomial Test p=0.634). However, we noted a decrease in the total frequency of recombinants 10-22 hours post-heat shock (Figure 2, Fisher’s Exact Test p=0.002), including crossover recombinant progeny. If the SCR assay was detecting SSA repair, ablation of *xpf-1* should result in a severe reduction of ‘crossover’ progeny without altering observed frequencies of noncrossover progeny. Therefore, the occurrence of crossover recombinants in the *xpf-1* mutant suggests SSA does not contribute to the detected SCR assay repair outcomes. This result is not surprising, as multiple *C. elegans* studies demonstrated that mutagenic DNA repair pathways, including SSA, are only frequently utilized for meiotic DSB repair in mutants where recombination is impeded [7,15,27–29]. We further demonstrate that the second recombinant product of intersister crossing-over can be detected via PCR of GFP-progeny from the SCR assay (Figure S2), and intersister crossovers were also cytologically observed in the accompanying publication [30], reinforcing that the crossover progeny we observe are derived from bona fide intersister crossovers. Overall, our data suggests that XPF-1 promotes meiotic sister chromatid repair specifically in late meiotic prophase I.

### Mechanisms of sister chromatid recombination

Recombination mechanisms can be inferred from gene conversion tracts, which are DNA sequence changes that arise from nonreciprocal exchanges during recombination repair with a polymorphic template. To reveal mechanisms of meiotic sister chromatid repair, we engineered polymorphisms in the two tandem GFP cassettes within the SCR assay, thereby enabling detection of conversion tracts from recombination between nonallelic GFP sequences (Figure 3A). Wild-type noncrossovers displayed tracts ranging from a single to multiple polymorphism conversions spanning 567bp of sequence (Figure 3B). In all of these tracts, the polymorphism most proximal to the site of Mos1 excision (12bp downstream) was always converted, indicating that recombination intermediates remain local to the site of DSB induction (Figure 3B). With this polymorphism density, we did not observe restoration tracts, which are unconverted polymorphisms flanked by conversion events indicative of multiple template engagement, heteroduplex DNA mismatch correction, or nucleotide excision of joint molecules during recombination [31–33] (Figure 3B). Although interhomolog conversion tracts in other organisms exhibit frequent joint molecule migration and strand switching [33,34], our results suggest sister chromatid noncrossover repair in *C. elegans* is likely processive and does not involve extensive migration from the DSB site.

**FIGURE 3.**
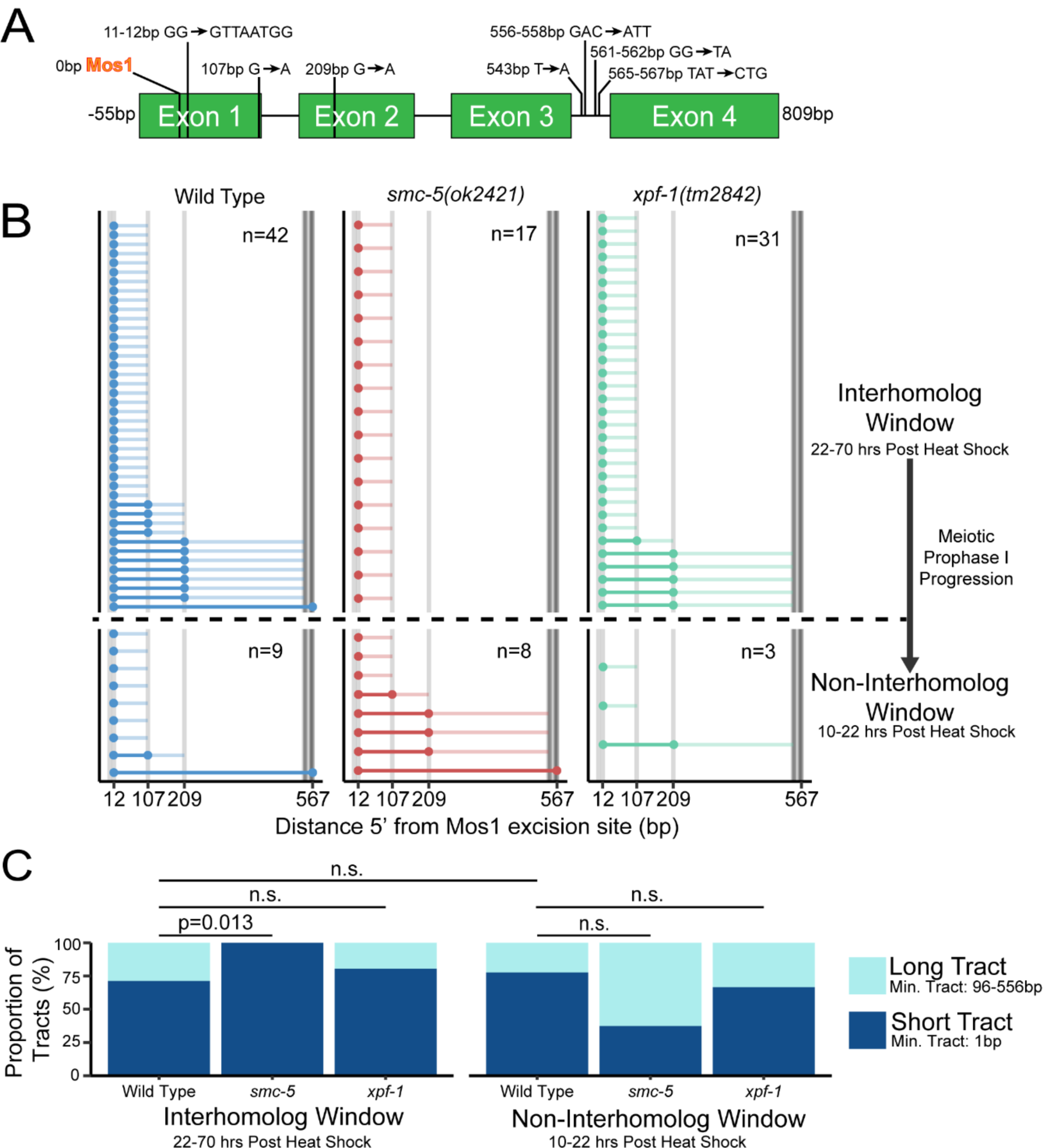
SMC-5 regulates sister chromatid noncrossover conversion tract length. (A) Scale cartoon of SCR assay GFP cassette with annotated polymorphisms. The polymorphisms of the *pmyo-2::*GFP sequence are listed to the left of each arrow, while the sequence of the *pmyo-3::GFP* polymorphism is listed to the right of each arrow. Positions of polymorphisms in bp are relative to the site of Mos1 excision. (B) Converted polymorphisms within SCR assay wild-type, *smc-5(ok2421)*, and *xpf-1(tm2842)* SCR assay noncrossover recombinant loci. Each horizontal line represents the sequenced locus of a single recombinant. High opacity lines connect contiguous converted polymorphisms within a single tract and represent minimum tract length, while the low opacity lines represent the range between converted and the most proximal non-converted polymorphism. (C) Stacked bar plots showing the proportion of ‘short’ (1bp minimum tract length) and ‘long’ (≥107bp minimum tract length). P values calculated by Fisher’s Exact Test.

To assess whether processing of sister chromatid recombination intermediates changes during meiotic progression, we compared tracts generated at different stages of prophase I (Figures 3B, 3C). The length of a conversion tract is a readout for the extent of 5’ strand resectioning, but may also be influenced by joint molecule migration, extent of strand synthesis, and mismatch repair of heteroduplex sequences [35,36]. Comparing the minimum conversion tract lengths of our wild-type noncrossover tracts, we note that the proportion of ‘short’ tracts converted only at one polymorphism and ‘long’ tracts 96-556 bp in length are similar within both the interhomolog and non-interhomolog windows (Figure 3C, Fisher’s Exact Test p>0.05), suggesting that early DSB processing during intersister and/or intrachromatid noncrossover recombination repair is likely similar throughout prophase I.

Notably, SCR assay noncrossover tracts share traits with interhomolog noncrossover tracts observed in other organisms. In mice, ∼71-84% of tracts at assayed hotspots are converted at only a single polymorphism [34,37], which is similar to the ∼72% of single polymorphism conversions in observed in our SCR assay (Figure 3B). Similarly, human NCO tracts are frequently singleton conversions [38,39]. This observation is divergent from *S. cerevisiae*, where meiotic noncrossover tracts are usually longer [40]. Unlike noncrossovers, crossovers in mice, humans, and *S. cerevisiae* display consistently larger tracts [34,37–40]. Similarly, we found that the smallest SCR assay intersister crossover tracts (198bp) are longer than their noncrossover counterparts (Figure S4). Although detection of conversion tracts varies significantly between organisms due to polymorphism density, the distribution of meiotic conversion tract sizes we observe in *C. elegans* using the SCR assay therefore appears concordant with other metazoans.

### Role of Smc5/6 in recombination mechanisms

To determine if SMC-5/6 influences DNA processing during DSB repair, we sequenced noncrossover tracts from *smc-5* mutant sister chromatid recombinants. Similar to wild-type, we observe consistent conversion of the most proximal polymorphism to the DSB in *smc-5* mutants (Figure 3B), suggesting that SMC-5/6 does not regulate joint molecule migration. However, we observed a significant absence of ‘long’ 96-556bp noncrossover tracts in the interhomolog window of *smc-5* mutants (Figure 3C, Fisher’s Exact Test p=0.013). These data reveal that although SMC-5/6 is required only in late meiotic prophase I for promoting sister chromatid repair, it also influences recombination intermediate processing during early-mid prophase I.

Several studies suggest Smc5/6 regulates DSB resection and joint molecule resolution. In *S. cerevisiae*, Smc5/6 promotes mitotic DSB resection by SUMOylating the STR complex [41,42]. In *C. elegans, smc-5* mutants exhibit elevated and persistent RAD-51 recombinase foci in pachytene [17]. Our observation of only short conversion tracts within the interhomolog window of *smc-5* mutants could indicate reduced resection, thereby impeding the homology search and prolonging the stage a DSB is detectable as a RAD-51 coated filament. In *C. elegans*, SMC-5/6 is also suggested to regulate chromatin structure in late meiotic prophase I to permit efficient DSB repair by preventing aberrant joint molecules [41]. In *S. cerevisiae*, Smc5/6 is required for recruiting the endonuclease Mus81 and the STR complex to promote cleavage of double Holliday junction and multi-chromatid joint molecule intermediates [18,42]. Our data find that SMC-5/6 is likely not required for resection in late prophase I but may regulate joint molecule resolution.

### Role of XPF-1 in recombination mechanisms

Similar to *smc-5* and wild-type, we observe no restoration tracts in *xpf-1* mutant sister chromatid noncrossovers, and the most DSB proximal polymorphism remained converted in every tract we sequenced (Figure 3B). The proportion of ‘short’ (1bp) and ‘long’ (95-556bp) noncrossover conversion tracts arising from *xpf-1* mutants were similarly indistinguishable from wild-type (Figure 3C, Fisher’s Exact Test interhomolog window and non-interhomolog window p>0.05). These results are consistent with XPF-1 functioning as a resolvase to promote sister chromatid repair in late meiotic prophase I following the DSB processing and joint molecule formation steps that influence conversion tract generation.

### *smc-5* and *xpf-1* mutations differentially affect progeny viability

The SCR assay directly assesses functions of specific proteins in sister chromatid repair but does not determine whether these functions affect gamete viability. To establish whether defects in sister chromatid recombination at specific stages of meiotic prophase I is required for fertility, we exposed young adult *smc-5* and *xpf-1* mutant hermaphrodites to ionizing radiation, which induces DSBs, and performed a reverse time-course to assess effects on brood viability of damage induced at specific meiotic stages. As previously reported [17], the overall cohort of *smc-5* mutant oocytes are more sensitive than wild-type to ionizing radiation throughout meiotic prophase I (Figure 4, Mann-Whitney U test p<0.001 at all timepoints). With our reverse time-course irradiation, we find that *smc-5* mutant oocytes (at 2500 and 5000 Rad) within the non-interhomolog window were hypersensitive to irradiation compared to the interhomolog window (Figure 4, Mann-Whitney U test p<0.001 for both treatments). Thus, SMC-5/6 is required specifically in late meiotic prophase I to ensure progeny viability, which corresponds to the stages when this complex promotes efficient sister chromatid repair and sister chromatid crossovers (Figures 2, 3).

**FIGURE 4.**
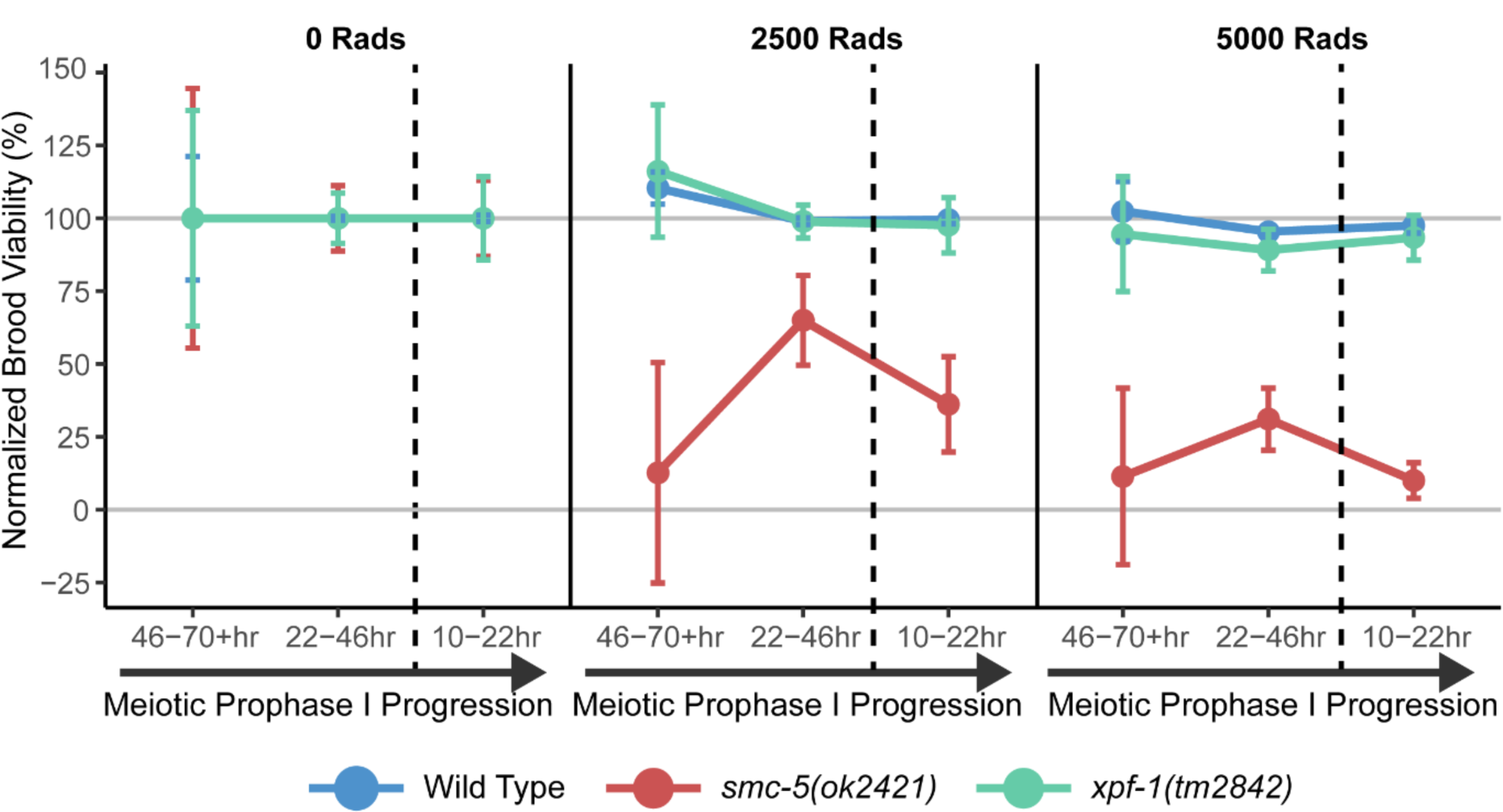
Mutant germline hypersensitivity to ionizing radiation is meiotic stage-dependent. Mean brood viability of young adult hermaphrodites exposed to 0, 2500, or 5000 Rads of ionizing radiation, normalized to the mean brood viability for each genotype and timepoint scored in the absence of ionizing radiation (0 Rads treatment). Broods of n=15 parent hermaphrodites of each respective genotype were scored for each irradiation treatment dose. Vertical dashed lines delineate between timepoints representing damage induced during the interhomolog window (22-70+ hrs) and timepoints representing damage induced during the non-interhomolog window (10-22 hrs). Error bars represent standard deviation. For pairwise statistical comparisons, see Supplemental Figure 3. Brood viabilities of each condition without normalization are displayed in Supplemental Figure 5.

Mutants for *xpf-1* exhibited a mild but significant reduction in brood viability upon exposure to 5000 Rads of ionizing radiation only within the 22-46 hour timepoint (Figure 4, Mann-Whitney U test p=0.027). While our SCR assay demonstrates that XPF-1 is required for efficient sister chromatid repair, *xpf-1* mutants were not radiation-sensitive in late meiotic prophase I. Therefore, XPF-1 might distinguish between DSBs generated from transposition versus ionizing irradiation. Alternately, this discrepancy in ionizing radiation sensitivity between *smc-5* and *xpf-1* mutants could reflect that defects in different steps of recombination do not equally impact progeny viability. A recent study found that *smc-6* mutants exhibited increased chromosomal structural variations following treatment with DNA damaging agents, while *xpf*-*1* exhibited less severe genetic lesions following irradiation [44]. Thus, mutants that affect intersister recombination late in joint molecule resolution are possibly less impactful on fertility and may result in DSB resolution by an error prone repair pathway. Mutants affecting earlier recombination steps may therefore be susceptible to catastrophic genomic rearrangements. Altogether, these data reinforce the meiotic-stage specific requirements of DNA repair complexes in maintaining genomic integrity.

## Conclusions

In this study we detected recombination between sister chromatids, thereby directly demonstrating that the sister chromatid is engaged as a repair template during meiosis. Further, we show that the Smc5/6 complex and XPF nuclease are differentially engaged to promote sister chromatid repair. We propose that SMC-5/6 performs multiple roles in promoting sister chromatid repair by influencing recombination intermediate processing in early prophase I and promoting efficient sister chromatid repair and intersister crossovers in late meiotic prophase I. We further demonstrate that the XPF-1 nuclease acts downstream of recombination intermediate processing to promote sister chromatid repair late in meiotic prophase I. Multiple repair pathways likely work with or in parallel to SMC-5/6 and XPF-1 to promote meiotic intersister recombination, and the SCR assay we have developed will enable future elucidation of these interactions.

## Acknowledgements

We thank the CGC (funded by National Institutes of Health (NIH) P40 OD010440) for strains. We thank A. Villeneuve, C. Cahoon, and N. Kurhanewicz for comments on the manuscript. We also thank O. Rog and D. Almanzar for sharing their manuscript and data prior to publication. This work was supported by the National Institutes of Health T32GM007413 to ET; National Institutes of Health R25HD0708 to CC, AH, and AS; and, National Institutes of Health R00HD076165 and R35GM128890 to DEL. DEL is also a recipient of a March of Dimes Basil O’Connor Starter Scholar award and Searle Scholar Award.

---

**SUPPLEMENTAL FIGURE 1.**
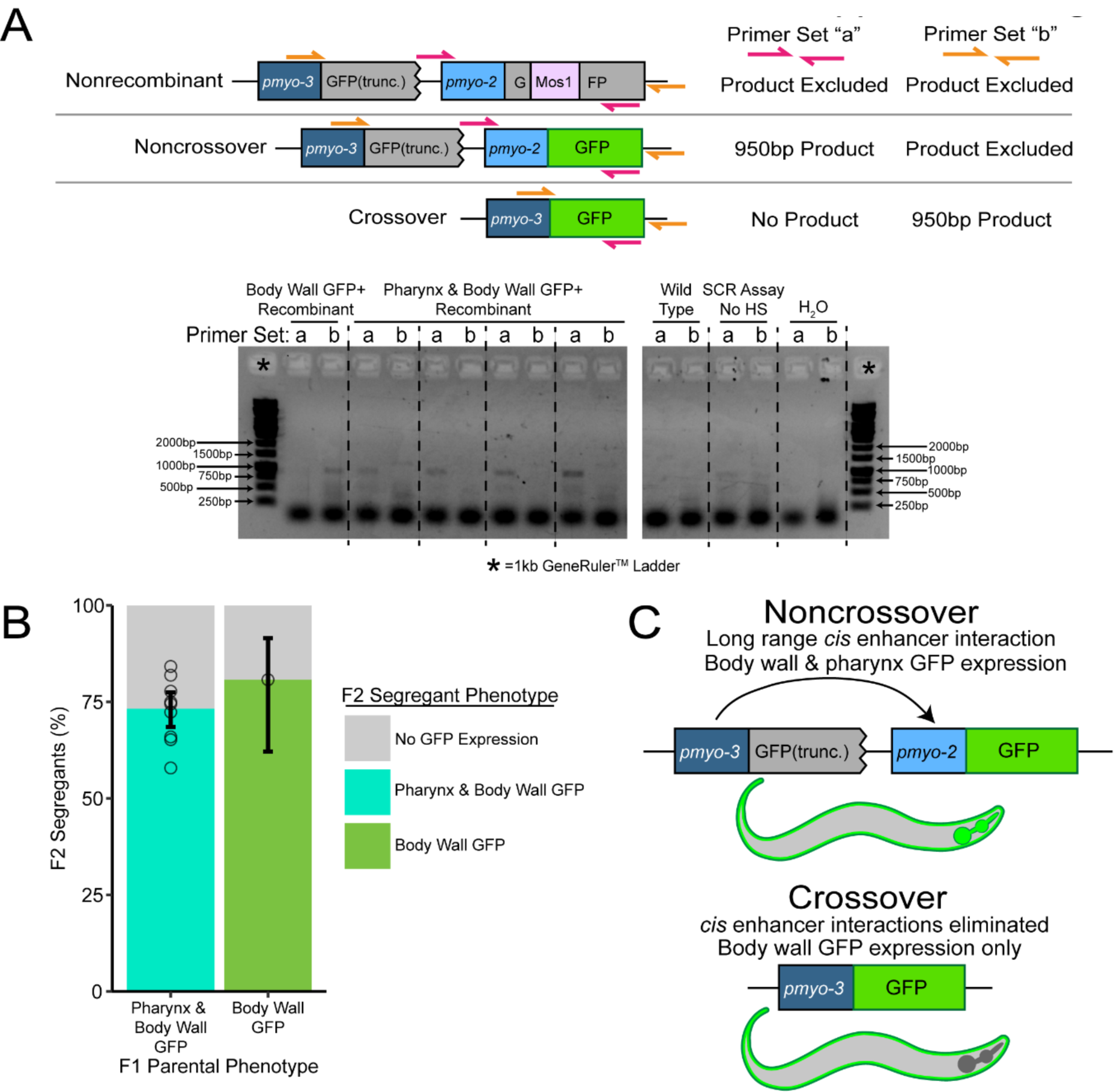
Sister chromatid repair assay recombinant progeny expressing GFP in multiple tissues are likely single noncrossovers. The design of the SCR assay predicts that recombinant progeny should exhibit either body wall or pharynx GFP expression (Figure 1A). However, the majority of pharynx GFP+ progeny also expressed GFP in the body wall. (A) PCR screening of recombinant progeny demonstrates expected crossover and noncrossover products in body wall and body wall + pharynx GFP expressing recombinants. ‘PCR product restricted’ indicates primer sets unable to amplify the given sequence based on the extension time of the PCR cycle. Wells in ethidium bromide stained agarose gel marked with asterisks were loaded with 1kb GeneRuler DNA ladder. (B) Both body wall and body wall + pharynx expression phenotypes segregate in ratios consistent with dominant mendelian traits arising from a single locus (Chi Square Test of Goodness of Fit p>>0.05 for both parental phenotypes). Error bars represent 95% Binomial confidence intervals, bars indicate proportion of segregants of each phenotype across all broods scored. Circles represent the proportion of segregants with respective phenotypes from the broods of individual F1 recombinant hermaphrodites scored. (C) Body wall + Pharynx GFP expression likely arises from known long-range enhancer activity between *myo-2* and *myo-3* promoter sequences [45].

**SUPPLEMENTAL FIGURE 2.**
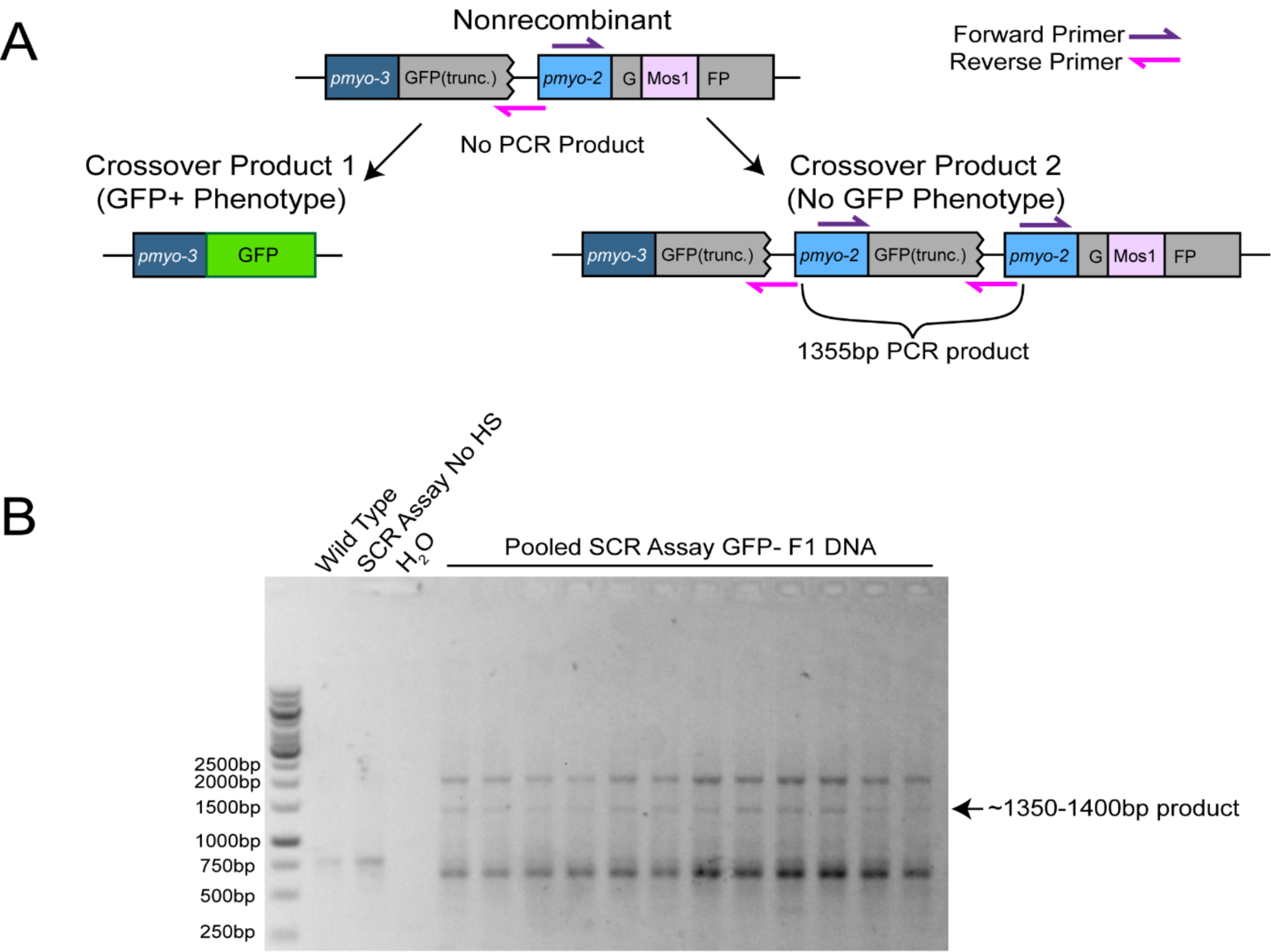
PCR detection of intersister crossover products in sister chromatid repair assay F1 progeny. (A) PCR design enabling selective detection of crossover recombinant products. Primers design ensures that a consistent product of 1355bp should only be generated from the non-GFP+ crossover product. However, as the primers bind multiple positions within the SCR assay and resultant recombinant constructs, it is likely that secondary priming would lead to additional off-target amplification. (B) An amplified PCR product consistent with the expected size for intersister crossover product 2 is detectable on an ethidium bromide stained agarose gel.

**SUPPLEMENTAL FIGURE 3.**
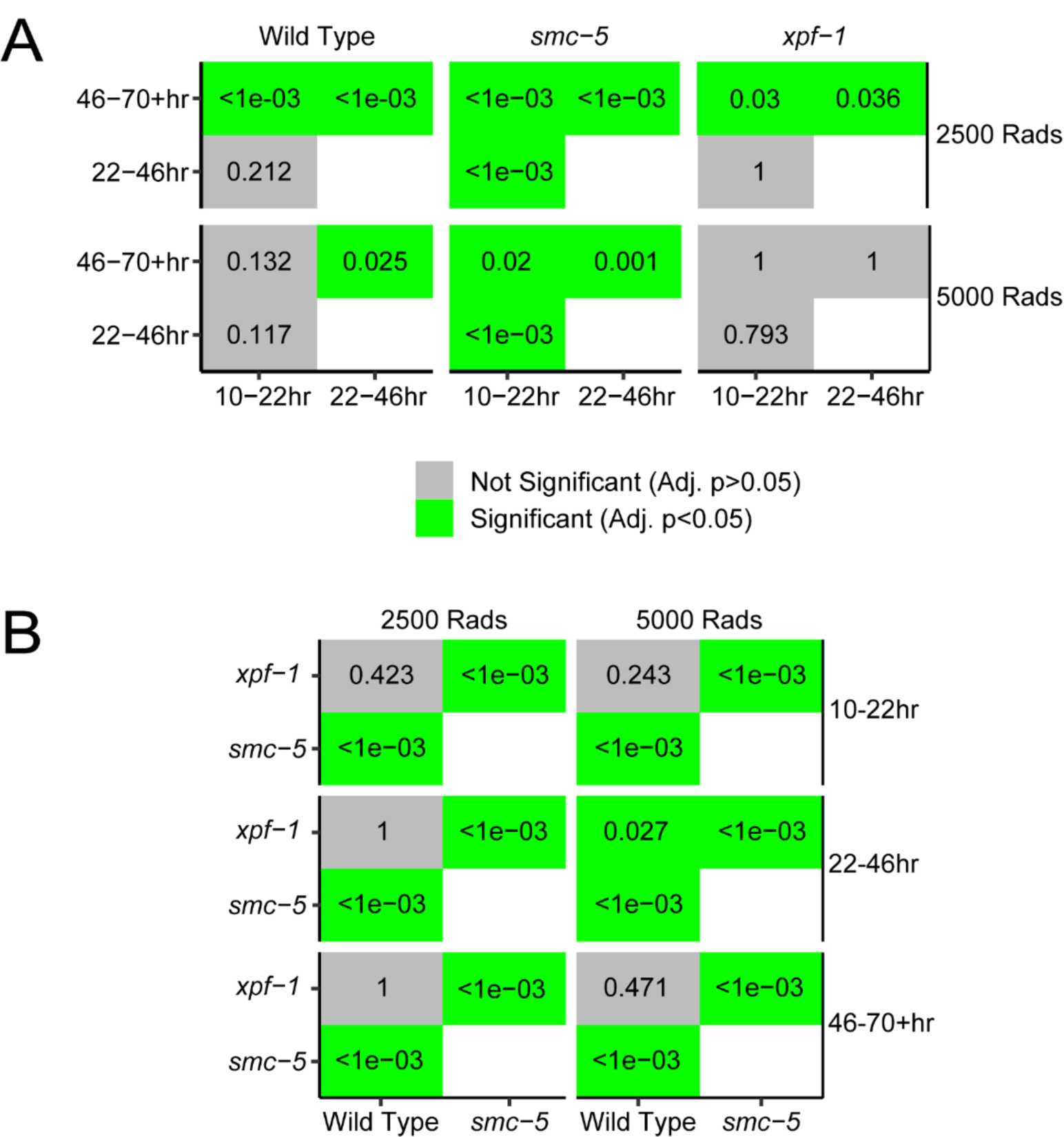
Pairwise statistical comparisons of brood viability data. All numbers on the plots indicate p values for respective comparisons from Mann Whitney U tests with Bonferroni corrections for multiple comparisons. Green boxes indicate significant values (p<0.05 after correction). (A) Comparisons of timepoints scored within the same genotype and treatment on raw brood viability data. (B) Comparisons for brood viability normalized to unirradiated control between indicated genotypes within the same treatment and timepoint scored.

**SUPPLEMENTAL FIGURE 4.**
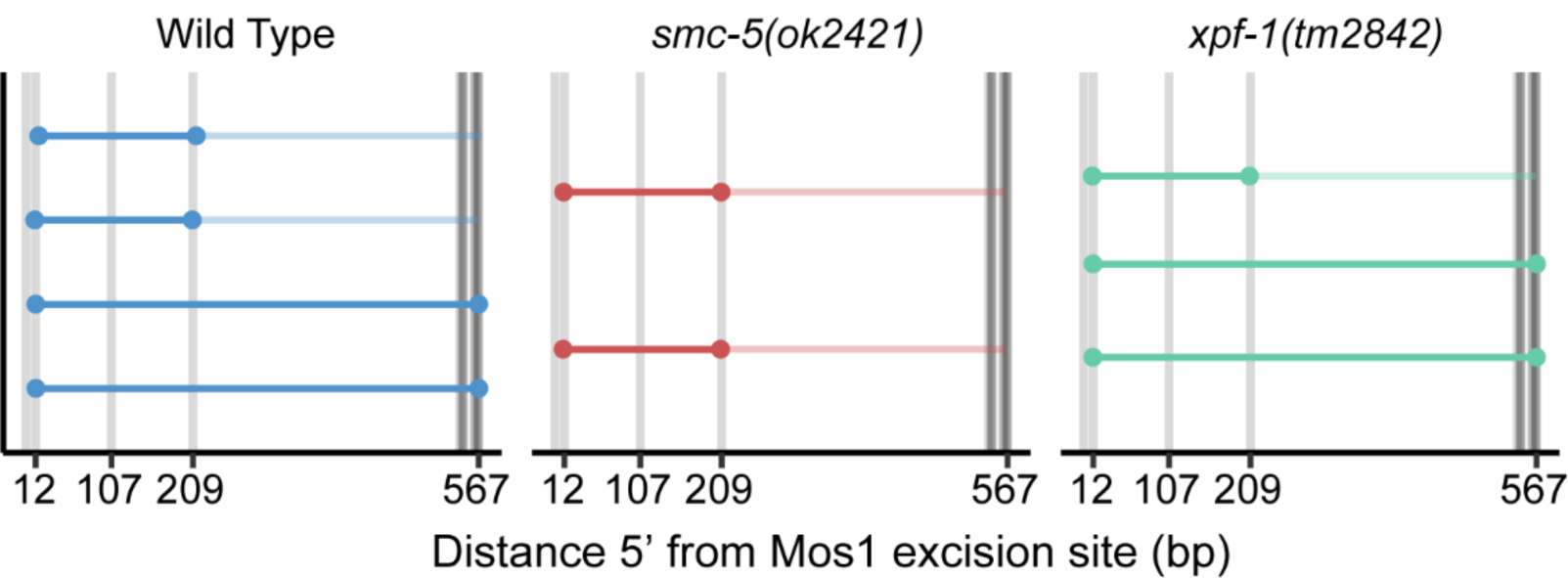
Intersister crossover conversion tracts. Converted polymorphisms within SCR assay wild-type, *smc-5(ok2421)*, and *xpf-1(tm2842)* SCR assay crossover recombinant loci. Each horizontal line represents the sequenced locus of a single recombinant. High opacity lines connect contiguous converted polymorphisms within a single tract and represent minimum tract length, while the low opacity lines represent the range between converted and the most proximal non-converted polymorphism.

**SUPPLEMENTAL FIGURE 5.**
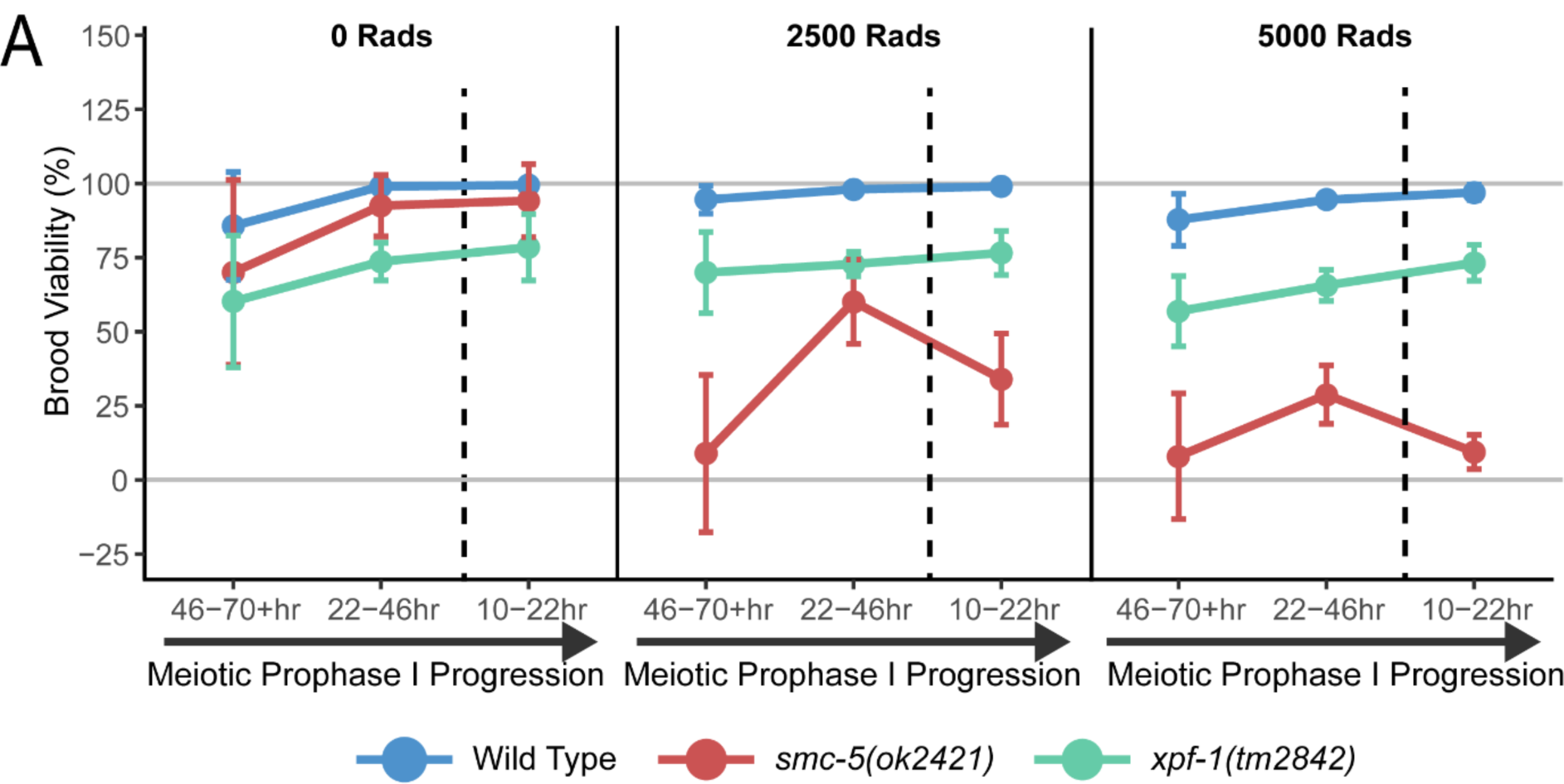
Mutant germline sensitivity to ionizing radiation. Mean brood viability of young adult hermaphrodites exposed to 0, 2500, or 5000 Rads of ionizing radiation. Broods of n=15 parent hermaphrodites of each respective genotype were scored for each irradiation treatment dose. Vertical dashed lines delineate between timepoints representing the interhomolog window (22-70+ hrs) and timepoints representing the non-interhomolog window (10-22hrs). Error bars represent standard deviations.

**Supplementary Table 1.** Recombination Frequency Counts of SCR Assay in Wild-type.

| Timepoint post Heat Shock | Total Progeny | Noncrossover | Crossover | % Recombinant Progeny (95% CI) | % Crossovers (95% CI) |
| --- | --- | --- | --- | --- | --- |
| 10-22 hours | 3317 | 19 | 7 | 0.78%<br>(0.54-1.15%) | 26.9%<br>(13.7-46.1%) |
| 22-34 hours | 2372 | 17 | 2 | 0.80%<br>(0.51-1.25%) | 10.5%<br>(2.9-31.4%) |
| 34-46 hours | 3032 | 16 | 1 | 0.56%<br>(0.35-0.90%) | 5.8%<br>(1.0-27.0%) |
| 46-58 hours | 2159 | 12 | 1 | 0.60%<br>(0.35-1.03%) | 7.7%<br>(1.4-33.3%) |
| 58-70 hours | 1190 | 8 | 1 | 0.76%<br>(0.40-1.43%) | 12.5%<br>(2.0-43.5%) |
| 70+ hours | 806 | 3 | 0 | 0.37%<br>(0.13-1.10%) | 0%<br>(0-56.1%) |
| Interhomolog Window (22-70+ hours) | 9559 | 56 | 5 | 0.64%<br>(0.50-0.82%) | 8.2%<br>(13.7-46.1%) |
| Total | 12876 | 75 | 12 | 0.68%<br>(0.55-0.83%) | 13.8%<br>(8.1-22.6%) |

**Supplementary Table 2.** Recombination Frequency Counts of Interhomolog Assay.

| Timepoint post Heat Shock | Total Progeny | Total Recombinant Progeny (Crossover:Noncrossover) | % Recombinant (95% CI) | % CO (95% CI) |
| --- | --- | --- | --- | --- |
| 10-22 hours | 1623 | 2 (0:2) | 0.12%<br>(0.03-0.45%) | 0%<br>(0-65.8%) |
| 22-34 hours | 1938 | 51 (3:38) | 2.56%<br>(1.96%-3.36%) | 7.3%<br>(2.5-19.4%) |
| 34-46 hours | 1629 | 92 (8:81) | 5.35%<br>(4.38-6.51%) | 8.9%<br>(4.6-16.7%) |
| 46-58 hours | 1369 | 78 (5:68) | 5.40%<br>(4.34%-6.68%) | 6.8%<br>(3.0-15.1%) |
| 58-70 hours | 691 | 19 (2:15) | 2.68%<br>(1.72%-4.14%) | 11.8%<br>(3.3-34.3%) |
| 70+ hours | 376 | 8 (0:8) | 2.08%<br>(1.06-4.06%) | 0%<br>(0-32.4%) |
| Interhomolog Window (22-70+ hours) | 6251 | 248 (18:210) | 3.97%<br>(3.51-4.48%) | 7.9%<br>(5.1-12.1%) |
| Total | 7876 | 250 (18:212) | 3.17%<br>(2.81-3.58%) | 7.8%<br>(5.4-13.0%) |

**Supplementary Table 3.**

Recombination Frequency Counts of *smc-5(ok2421)* SCR Assay
| Timepoint post Heat Shock | Total Progeny | Noncrossover | Crossover | % Recombinant (95% CI) | % CO (95% CI) |
| --- | --- | --- | --- | --- | --- |
| 10-22 hours | 3491 | 11 | 0 | 0.32%<br>(0.2-0.59%) | 0%<br>(0-25.9) |
| 22-34 hours | 2188 | 15 | 0 | 0.69%<br>(0.42-1.13%) | 0%<br>(0-20.4) |
| 34-46 hours | 1009 | 2 | 1 | 0.30%<br>(0.10-0.87%) | 33%<br>(6.1-79.2) |
| 46-58 hours | 433 | 3 | 1 | 0.92%<br>(0.35-2.35%) | 25%<br>(4.6-69.9) |
| 58-70 hours | 121 | 0 | 0 | 0%<br>(0-3.8%) | -- |
| 70+ hours | 72 | 0 | 0 | 0%<br>(6.94e-18-5.1%) | -- |
| Interhomolog Window (22-70+ hours) | 3801 | 20 | 2 | 0.58%<br>(0.38-0.87%) | 9.1%<br>(2.5-27.8%) |
| Total | 7314 | 31 | 2 | 0.45%<br>(0.32-0.63%) | 6.1%<br>(1.7-19.6%) |

Recombination Frequency Counts of *xpf-1(tm2842)* SCR Assay
| Timepoint post Heat Shock | Total Progeny | Noncrossover | Crossover | % Recombinant (95% CI) | % CO (95% CI) |
| --- | --- | --- | --- | --- | --- |
| 10-22 hours | 2618 | 4 | 2 | 0.23%<br>(0.11-0.50%) | 33.3%<br>(9.7-70%) |
| 22-34 hours | 1793 | 9 | 2 | 0.61%<br>(0.34-1.1%) | 18.2%<br>(5.1-47.7%) |
| 34-46 hours | 2400 | 9 | 2 | 0.46%<br>(0.26-0.82%) | 18.2%<br>(5.1-47.7%) |
| 46-58 hours | 1819 | 11 | 0 | 0.61%<br>(0.39-1.1%) | 0%<br>(0-25.9%) |
| 58-70 hours | 813 | 4 | 0 | 0.49%<br>(0.19-0.13%) | 0%<br>(0-49.0%) |
| 70+ hours | 469 | 1 | 0 | 0.21%<br>(0.04-1.2%) | 0%<br>(0-79.3%) |
| Interhomolog Window (22-70+ hours) | 7294 | 34 | 4 | 0.52%<br>(0.38-0.71%) | 10.5%<br>(4.2-24.1%) |
| Total | 9912 | 38 | 6 | 0.44%<br>(0.33-0.60%) | 13.6%<br>(6.4-26.7%) |

## STAR METHODS

### LEAD CONTACT AND MATERIALS AVAILABILITY

Requests for further information, reagents, or resources should be directed to the lead contact, Diana E. Libuda.

### EXPERIMENTAL MODEL AND SUBJECT DETAILS

#### *C*. *elegans* Strain Maintenance and Strain Construction

*C. elegans* strains used in this study were maintained at 15°C or 20°C on nematode growth medium (NGM) plates and were fed the OP50 *Escherichia coli* bacterial strain. Experiments were performed only on *C. elegans* strains that had been maintained at 20°C for a minimum of two generations.

*C. elegans* strains used in this study were obtained from the *Caenorhabditis* Genetics Center (CGC) or were generated by crossing and/or CRISPR/Cas9 genome editing. Genetic crosses were performed by placing L4 stage male and hermaphrodite nematodes on NGM plates with OP50 at 20°C and screening for cross progeny after 3-4 days. Genotypes of strains generated by crossing were confirmed by PCR. DLW23 was generated by crossing YE57 males to DLW14 hermaphrodites. To generate DLW82, TG1660 hermaphrodites were crossed to YE57 males to generate males carrying the *xpf-1(tm2842)* allele balanced by the mIn1 balancer. These F1 *xpf-1*/mIn1 males were then crossed to DLW14.

Strains generated by CRISPR/Cas9 genome editing were backcrossed to remove any off-target mutations that may have been incurred. The strain carrying the integrated sister chromatid repair assay sequence, DLW14, was backcrossed three times to EN909.

### METHOD DETAILS

#### Sister Chromatid Repair Assay Construction

The sister chromatid repair assay plasmid pMG1 was constructed by integrating *pmyo-3* sequence from pCFJ104 (Jorgensen Lab) into the synthetic plasmid pDL23 (GenScript) by Gibson assembly (SGI-DNA) and PCR stitching. Plasmids pMG3 and pMG14 expressing Cas9 and CRISPR guide RNAs (pMG3 protospacer 5’-GAGUAGUUCAGGAUCUGG-3’, pMG14 protospacer 5’-GUUGUUGAAUGUGGUAGAGG-3’) targeting *unc-5* were generated by modifying pJW1285 (Jorgensen Lab) using PCR stitching. All plasmid sequences were confirmed by Sanger sequencing (Sequetech).

#### CRISPR/Cas9 *C*. *elegans* Genome Editing

CRISPR/Cas9 genome editing to integrate the sister chromatid repair assay into the *unc-5* locus on Chromosome IV of the *C. elegans* genome was performed by injecting the germlines of adult N2 hermaphrodites with a plasmid mix (100ng/μL pMG1, 30ng/μL pMG3, 30ng/μL pMG14). F1 progeny of injected hermaphrodites were screened for uncoordinated movement (Unc) phenotypes, indicating editing at the *unc-5* locus. Integration of the SCR assay was confirmed by PCR, and the entire integrant construct sequence was confirmed by Sanger sequencing (Sequetech).

#### Sister Chromatid Repair Assay

Parent (P0) hermaphrodites for sister chromatid repair assays were generated by crossing. L4 stage P0 hermaphrodites were picked 16-18 hours before heat shock and incubated overnight at 15°C. To improve progeny yields at later timepoints, where the abundance of hermaphrodite sperm limits brood size, N2 young adult males were added to these plates in some replicates of the SCR assay. Heat shock was performed by placing P0 hermaphrodites in an air incubator (refrigerated Peltier incubator, VWR Model VR16P) at 34°C for one hour. Following heat shock, hermaphrodites were incubated at 20°C for 10 hours and then were picked to individual NGM plates seeded with OP50. After 12 hours each P0 hermaphrodite was transferred to a new NGM plate. P0 hermaphrodites were similarly passaged to new NGM plates every 12 hours for a total of 6 transfers. NGM plates with P0 hermaphrodites were maintained at 20°C, while NGM plates with F1 progeny only were placed at 15°C.

F1 progeny were maintained at 15°C for 36-48 hours. ∼18 hours before scoring for fluorescence, F1 progeny were placed in a 25°C incubator to enhance GFP expression. F1 progeny were scored for fluorescence using an Axio Zoom V16 fluorescent dissection microscope (Zeiss). F1s that expressed GFP in the pharynx, body wall, or both were transferred to individual plates for single worm lysis (as described in Sister Chromatid Repair Assay Conversion Tract Analysis methods). All other progeny were removed from the plate and discarded. If all F1 progeny were in larval developmental stages at the time of scoring, dead eggs and unfertilized oocytes on the plates were additionally quantified.

We noted that the majority of recombinant progeny with *pmyo-2* (pharynx) GFP expression also exhibited *pmyo-3* (body wall) GFP fluorescence. To determine if this expression pattern arose from a single locus, we assayed the segregation of GFP phenotypes in F2 progeny arising from pharynx and body wall GFP expressing SCR assay progeny. The ratios of segregation were consistent with Mendelian inheritance of a single locus (Supplemental Figure 1B). PCR genotyping of progeny with both *pmyo-2* (pharynx) and *pmyo-3* (body wall) GFP expression produced products consistent with the presence of noncrossover recombination events specifically (Supplemental Figure 1A). Previous work has demonstrated that both the *pmyo-2* and *pmyo-3* promoters contain enhancers that alter the specificity of the other respective promoter’s expression pattern [45]. We therefore suggest that recombinants with both pharynx and body wall GFP expression patterns arise from the enhancer activity of the upstream *myo-3* promoter in noncrossover recombinants (Supplemental Figure 1C). Progeny exhibiting both pharynx and body-wall GFP expression were scored as noncrossover recombinants in all recombination frequency calculations.

We additionally found that a fraction of F1 SCR assay progeny exhibited weak fluorescence phenotypes only in a portion of the pharynx, body wall, or both tissues. These progeny were transferred to individual plates and maintained at 20°C. F2 progeny were visually screened for inheritance of a fluorescent phenotype. No partial tissue fluorescent F1 was ever observed to produce fluorescent progeny, indicating that these fluorescent phenotypes are a product of somatic Mos1 excision and subsequent DNA repair and are not the result of bona fide meiotic recombination. Partially fluorescent F1s were categorized as nonrecombinant when determining frequencies of meiotic sister chromatid recombination.

The sister chromatid repair assay was replicated a minimum of three times for genotype.

While performing sister chromatid repair assays in N2 and *xpf-1(tm2842)* backgrounds in which the *unc-5(lib1)* and KrIs14 transgenes were inherited from a hermaphrodite, we observed a spontaneous change in results encompassing: (1) reduced recombinants at the 10-22 hour timepoint following heat shock; and, (2) severe embryonic lethality amongst progeny laid 22+ hours following heat shock. We were able to successfully restore function of the SCR assay by performing cross schemes to ensure that the parent hermaphrodites heat shocked in the SCR assay inherited their *unc-5(lib1)* allele and KrIs14 transgene from a male. We therefore recommend that future SCR assays only be performed on parent hermaphrodites who inherit these transgenes from a male. For descriptions of both cross schemes, see ‘Crosses to Generate Strains for the Sister Chromatid Repair Assay’.

#### Crosses to Generate Strains to Perform the Sister Chromatid Repair Assay

1. N2 (wild-type) with SCR assay transgenes inherited from hermaphrodite: Parent hermaphrodites were generated by crossing: (1) DLW14 hermaphrodites x N2 males to generate F1 *unc-5(lib1)*/*+* IV; KrIs14/+ V hermaphrodites.
2. Cross scheme 2:N2 (wild-type) with SCR assay transgenes inherited from male: Parent hermaphrodites were generated by crossing: (1) N2 males x DLW14 hermaphrodites to generate F1 *unc-5(lib1)*/*+* IV; KrIs14/+ V males, (2) F1 males x CB791 hermaphrodites to generate *unc-5(lib1)*/*unc-5(e791)* IV; KrIs14/+ V hermaphrodites.
3. *smc-5* with SCR assay transgenes inherited from hermaphrodite: Parent hermaphrodites were generated by crossing: (1) YE57 males x DLW23 hermaphrodites to generate *smc-5(ok2421)*/*smc-5(ok2421)* II; *unc-5(lib1)*/*+* IV; KrIs14/+ V hermaphrodites.
4. *xpf-1* with SCR assay transgenes inherited from male: Parent hermaphrodites were generated by crossing: (1) YE57 males x TG1660 hermaphrodites to generate *xpf-1(tm2842)*/mIn1 II males, (2) F1 males x DLW75 hermaphrodites to generate *xpf-1(tm2842)*/mIn1 II; *unc-5(lib1)*/*+* IV; KrIs14/+ V males, (3) F2 males x DLW82 hermaphrodites to generate *xpf-1(tm2842)*/ *xpf-1(tm2842)* II; *unc-5(lib1)*/*unc-5(e791)* IV; KrIs14/+ V hermaphrodites.
5. *xpf-1* with SCR assay transgenes inherited from male: Parent hermaphrodites were generated by crossing: (1) TG1660 males x DLW75 hermaphrodites to generate *xpf-1(tm2842)*/mIn1 II; *unc-5(lib1)*/*+* IV; KrIs14/+ V males, (2) F1 males x DLW82 hermaphrodites to generate *xpf-1(tm2842)*/ *xpf-1(tm2842)* II; *unc-5(lib1)*/*unc-5(e791)* IV; KrIs14/+ V hermaphrodites.

#### Sister Chromatid Repair Assay Conversion Tract Analyses

Genomes of fluorescent recombinant F1 progeny or the fluorescent F2 segregants of isolated recombinant F1 progeny from sister chromatid repair assays were extracted by single worm lysis. Individual hermaphrodites were picked into single 10μL aliquots of worm lysis buffer (50mM KCl, 10mM TrisHCl pH 8.2, 2.5mM MgCl_2_, 0.45% IGEPAL, 0.45% Tween20, 0.3μg/μL proteinase K in ddH_2_O). Each suspended worm was then serially frozen and thawed three times by immersion in a 95% ethanol and dry ice bath followed by a 65°C water bath. Each lysate was incubated at 60°C for one hour and then incubated at 95°C for 15 minutes to heat inactivate proteinase K. Final lysates were diluted with 10μL of ddH_2_O.

Recombinant loci were PCR amplified using OneTaq 2x Master Mix (New England Biolabs). Specificity of PCR reactions was determined by gel electrophoresis. Desired amplicons were extracted by PCR purification (Zymo PCR Purification Kit) if only one band was observed by electrophoresis, or gel extraction (Thermo Scientific Gel Extraction Kit) if multiple amplicons were observed. Purified amplicons were submitted for Sanger sequencing (Sequetech) with sequencing primers specific to the locus (see primer table). Sequencing files were aligned to reference GFP sequences with Benchling alignment software to detect converted polymorphisms.

The most efficient and effective primer set for amplifying *pmyo-2* (pharynx) GFP+ loci was DLO822 + DLO823. In addition, *pmyo-2* (pharynx) GFP+ loci were amplified using DLO640 + DLO641. The most efficient and effective primer set for amplifying crossover loci was DLO824+DLO546.

Not all fluorescent progeny lysed were able to be PCR amplified or successfully sequenced. We were able to completely sequence 51/74 wild-type NCO recombinants, 4/13 wild-type CO recombinants, 25/32 *smc-5(ok2421)* NCO recombinants, 2/2 *smc-5(ok2421)* CO recombinants, 34/39 *xpf-1(tm2842)* NCO recombinants, and 3/4 *xpf-1(tm2842)* CO recombinants.

#### Interhomolog Assay

The interhomolog assay was replicated following the protocol outlined in Rosu, Libuda, and Villleneuve *Science* 2011. In brief, parent (P0) hermaphrodites for interhomolog assays were generated by crossing AV554 males to CB791 hermaphrodites to generate *dpy-13(e184sd) unc-5(ox171::Mos1)/+ unc-5(e791)* IV; KrIs14*/+* V F1 progeny. Heat shock was performed by placing P0 hermaphrodites in an air incubator (refrigerated Peltier incubator, VWR Model VR16P) at 34°C for one hour. Following heat shock, hermaphrodites were incubated at 20°C for 10 hours and then were picked to individual NGM plates seeded with OP50. After 12 hours each P0 hermaphrodite was transferred to a new NGM plate. P0 hermaphrodites were similarly passaged to new NGM plates every 12 hours for a total of 6 transfers. Following transfer, the number of eggs laid by each hermaphrodite was scored. ∼48-60 hours following transfer, F1 progeny were scored for recombinant phenotypes. For details in determining noncrossover and crossover progeny, see Rosu, Libuda, and Villeneuve *Science* 2011.

#### Ionizing radiation and quantification of brood viability

L4 stage hermaphrodites were picked 16-18 hours before irradiation and incubated overnight at 15°C. Irradiation was performed using a ^137^Cs source (University of Oregon). Following irradiation, hermaphrodites were singled to individual NGM plates with OP50 lawns and were maintained at 20°C. At 10hrs and 46hrs following irradiation, the hermaphrodites were transferred to new NGM plates seeded with OP50. At ∼70hrs post irradiation, irradiated parent hermaphrodites were discarded. The proportion of hatched F1 progeny, dead eggs, and unfertilized oocytes were scored 36-48 hours following hermaphrodite removal. Brood viability was calculated as (Hatched Progeny) / (Hatched Progeny + Dead Eggs). Normalized brood viability was calculated by dividing the brood viability of each irradiated hermaphrodite within each scored timepoint (10-22 hrs, 22-46 hrs, 46-70+ hrs) by the mean brood viability of unirradiated hermaphrodites. Brood viability experiments were replicated three times for each genotype and irradiation dose, with the broods of n=5 hermaphrodites scored per replicate.

#### Extraction of pooled SCR assay F1 GFP-segregant genomes

Following the protocols described above, the sister chromatid repair assay was performed using >30 parent hermaphrodites generated through cross scheme #2. However, instead of performing the full time course, parent hermaphrodites were discarded following the 22-34hr timepoint. GFP+ meiotic recombinant F1 progeny were removed from the plates, and remaining F1 ‘GFP-’ progeny were washed from plates with worm lysis buffer without proteinase K (50mM KCl, 10mM TrisHCl pH 8.2, 2.5mM MgCl_2_, 0.45% IGEPAL, 0.45% Tween20 in ddH_2_O) and were frozen at −20°C for storage. DNA extraction was performed using a DNeasy Blood and Tissue Kit (Qiagen) following a modified version of the Kaganovich Lab Genomic DNA Isolation using Qiagen kit protocol. Upon thawing, worms were centrifuged at 2500 g and excess lysis buffer was removed. The pelleted progeny were resuspended in 200uL ATL buffer (DNeasy kit) and were freeze thawed 3x using liquid Nitrogen and a 65°C water bath. 20uL of Proteinase K (New England Biolabs) was added and the lysed worm solution was incubated at 56°C for 2 hours. 8uL of RNAse A (Sigma Aldrich) was added and the solution was incubated at room temperature for 5 minutes. 200uL of AL buffer (DNeasy kit) and the solution was incubated for 10 minutes at 56°C. 200uL of 100% ethanol was then added, and the solution was vortexed. Remaining steps of the protocol followed the published Qiagen kit instructions. All DNA was eluted from a single column in 50uL of ddH_2_O and was stored at −20°C until used.

#### Quantification of SCR assay F2 segregant phenotypes

Fluorescent recombinant SCR assay progeny were identified following the protocols described above using a total of 28 parent hermaphrodites generated through cross scheme #2. However, instead of performing the full time course, parent hermaphrodites were discarded following the 22-34hr timepoint. n=11 F1 progeny were identified that expressed GFP both in the body wall and in the pharynx and n=1 F1 progeny was identified that expressed GFP in the body wall only. Each of these recombinants was placed on an individual NGM plate seeded with OP50 and was incubated at 20°C. Each recombinant was monitored daily to determine if it had laid eggs. If >30 eggs were visible on the plate or the F1 recombinant was visually egg laying defective, identified by internal egg hatching inside of the F1, the F1 recombinant was lysed for PCR analysis (Figure S1a). F2 segregants were maintained at 20°C for an additional 24 hours, and then were scored for fluorescent phenotypes using an Axio Zoom V16 fluorescent dissection microscope (Zeiss).

### QUANTIFICATION AND STATISTICAL ANALYSIS

#### Statistics

Proportions of recombinant sister chromatid repair or interhomolog repair assay progeny and proportions of ‘short’ and ‘long’ conversion tracts (Figure 1b, Figure 2, Figure 3c) were compared using Fisher’s Exact Test. Determination of whether the number of observed intersister crossover recombinants in *smc-5* and *xpf-1* mutants deviated from expected wild-type frequencies was performed by Exact Binomial tests. Brood viability between timepoints within the same genotype and between genotypes within the same timepoints were compared using Mann-Whitney U tests with Bonferroni p value correction for multiple comparisons (Figure 4, Figure S3). Segregation ratios of F2 progeny from F1 SCR assay recombinants (Figure S1b) were compared to an expected distribution for mendelian segregation of a dominant phenotype arising from a single locus (75% parental phenotype, 25% no GFP expression) by Chi Square Tests of Goodness of Fit. For all tests, statistical significance was determined as a p value equal to or less than 0.5 following correction for multiple comparisons, if applicable. All statistical calculations were performed using R. 95% confidence intervals (Figure 1b, Figure 2, Figure 3c, Figure S1b) were calculated using the DescTools package in R.

### KEY RESOURCES TABLE

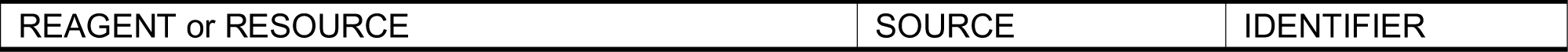
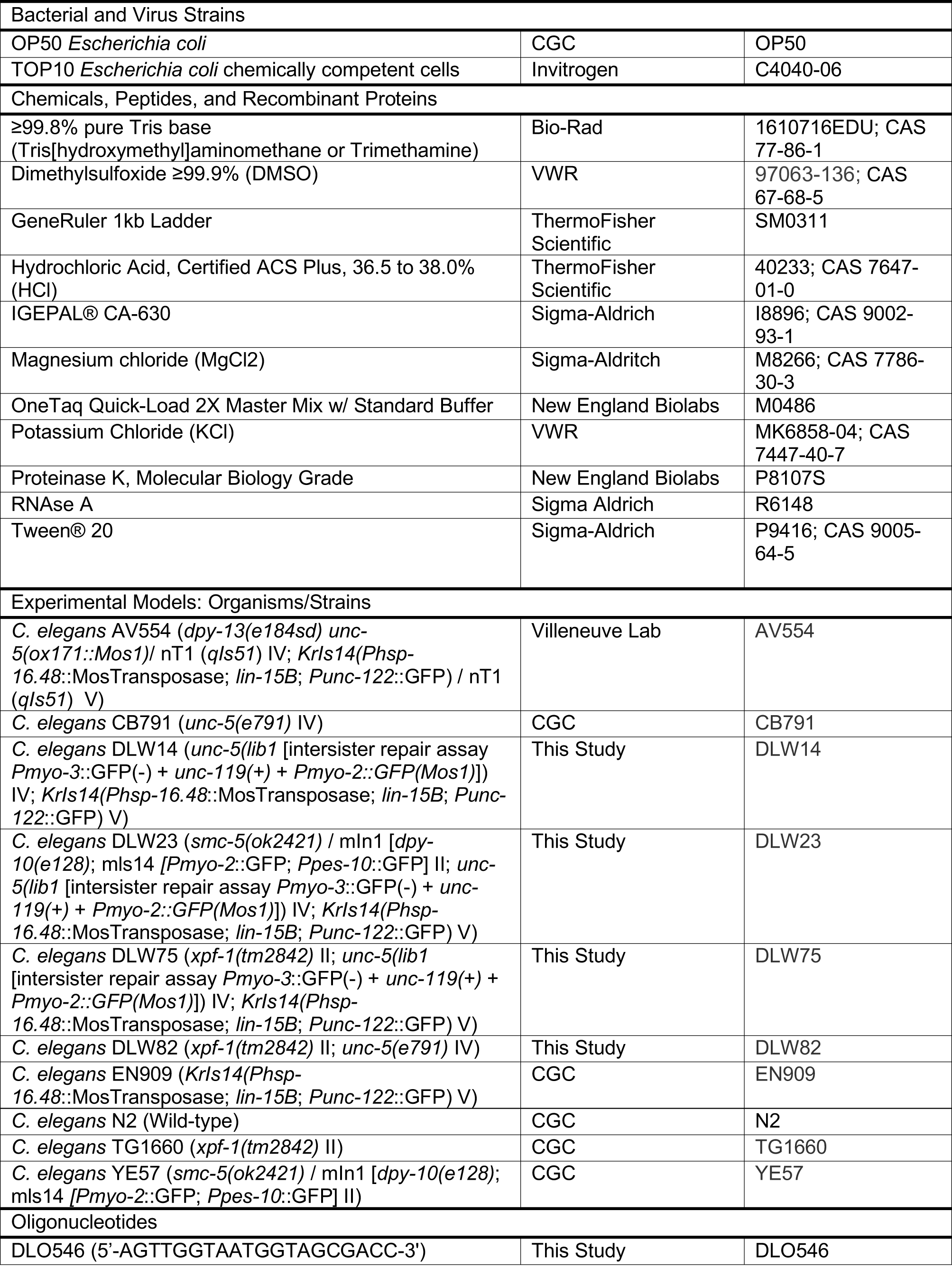
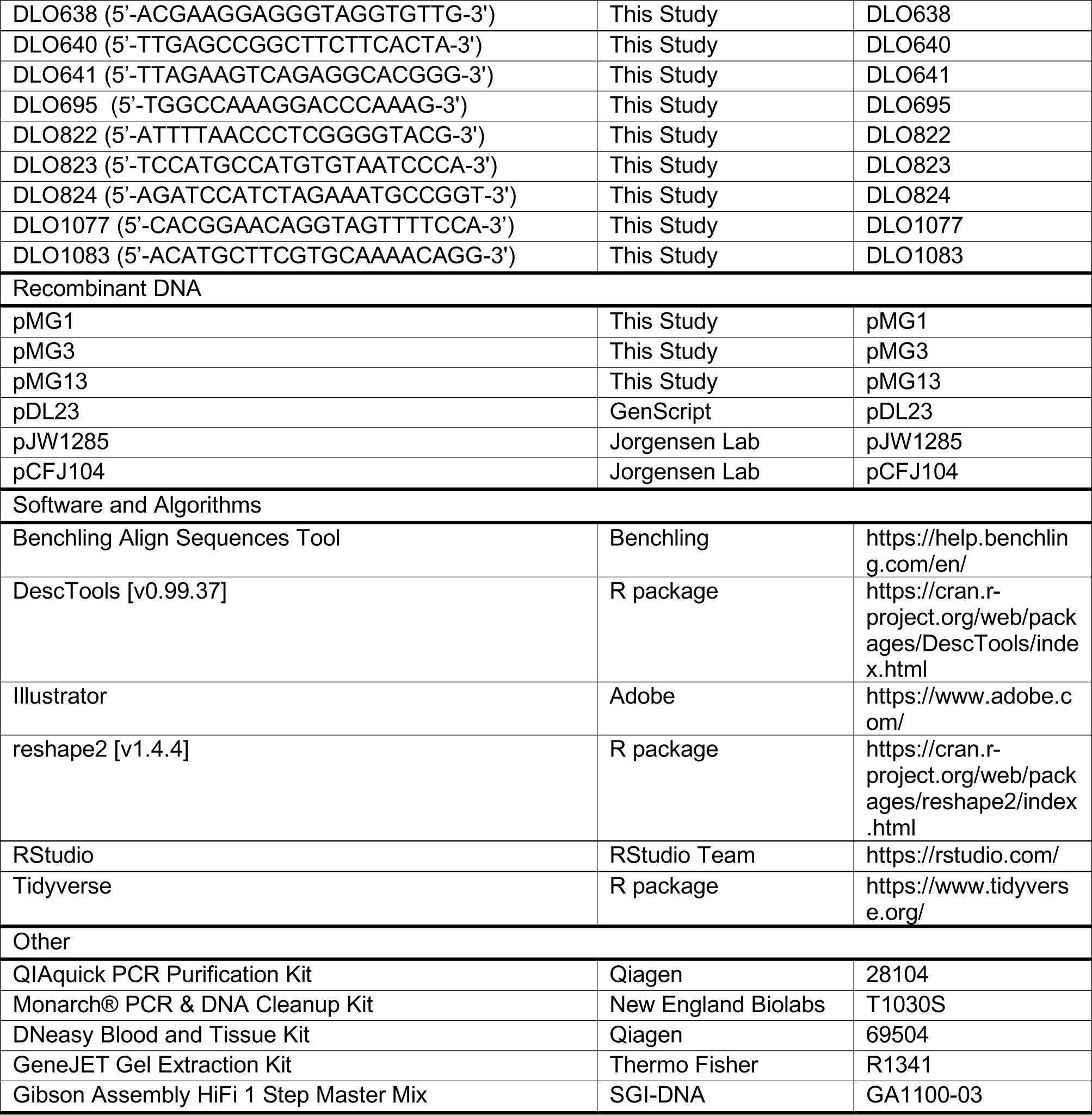

## Notes

### Competing Interest Statement

The authors have declared no competing interest.

### Summary of Updates

Figure 2 revised - colors in key for smc-5 mutant were swapped in previous version

